# A Multiresolution Hierarchical Approach to Peak Picking for High Resolution Mass Spectrometry Data Analysis

**DOI:** 10.64898/2025.12.07.692807

**Authors:** Vojtěch Bartoň, Ivana Ihnatova, Jiří Kalina, Garry Paul Codling, Elliott James Price, Helena Vítková, Vlad Popovici

## Abstract

**Motivation:** Accurate peak detection is a critical first step in high-resolution mass spectrometry (HRMS) data analysis. Most existing tools rely on centroiding and grid-based assumptions, which simplify the raw profile data at the cost of information loss and reduced detection accuracy.

**Results:** We present MRH, a novel peak detection method that operates directly on raw HRMS data using a hierarchical, multi-resolution decomposition of the (RT, *m/z*) space. Peaks are identified within resolution-dependent regions of interest using image-processing techniques, and an ambiguity score is introduced to quantify detection confidence. Benchmarking against widely used methods (apLCMS, centWave, and gridmass) shows that MRH consistently achieves higher F1 scores across concentrations, particularly excelling at low concentrations where competing tools fail. MRH also provides accurate localization of peaks and low ambiguity in detections, highlighting both robustness and precision.

**Availability:** A proof-of-concept implementation of MRH and the evaluation dataset are freely available at https://github.com/VojtechBarton/MRH.

## Introduction

High-resolution mass spectrometry (HRMS) is widely used for the identification of known and unknown compounds based on their mass-to-charge ratio (*m/z*). In untargeted screening, HRMS produces large datasets comprising spectra across retention times, which can be viewed as a two-dimensional function of intensity over (RT, *m/z*). Peak detection—the identification of significant local maxima in this signal—is a critical first step in downstream analysis and compound identification.

Despite its importance, defining what constitutes a “significant” peak remains challenging. Ideally, peaks correspond to chromatographic elution profiles with narrow *m/z* distributions and Gaussian-like retention time shapes, but real data often deviate from these assumptions. Moreover, the notion of locality in peak detection is complicated by discretization: while lower-resolution data may appear well aligned on a grid, higher-resolution data reveal gaps and misalignments, making simple Euclidean neighborhoods inadequate. Thus, multi-resolution approaches are better suited to capture data structure while avoiding prohibitive grid sparsity.

A further difficulty lies in data reduction. Many established tools (e.g. CentWave [1], MatchedFilter [1], ADAP-GC [2], mzMine2 [3], GridMass [4], XCMS [5]) rely on centroiding to simplify raw profile data. While centroiding reduces computational complexity, it can distort signals and lead to information loss that propagates through downstream analyses [6, 7]. Selecting centroiding parameters is especially problematic in untargeted studies, where prior knowledge of compound classes is limited.

Alternative approaches include direct processing of raw data and, more recently, the application of machine learning and deep learning [8, 9]. While promising, such methods require large annotated datasets and clear peak definitions—both of which remain difficult to obtain.

Here, we introduce MRH, a novel scalable method for peak detection that operates directly on raw HRMS data. The method decomposes data hierarchically into nested resolution levels, partitioning it into non-overlapping regions of interest that can be processed independently. This approach preserves maximum measurement precision, facilitates parallelization, and provides a flexible representation for downstream tasks. In addition, MRH introduces an ambiguity score, a simple metric for assessing the reliability of individual detections. Together, these innovations address key limitations of existing tools and provide a robust framework for peak detection in high-resolution metabolomics workflows.

## Methods

### Data representation and preprocessing

HRMS data are inherently irregularly sampled, particularly in the *m/z* domain, where measurements may not align to a regular lattice. At lower resolutions, points may appear aligned, but higher-resolution data reveal gaps and misalignments. This makes Euclidean distance on a fixed grid inadequate for defining neighborhoods. A multi-resolution representation alleviates this issue by adapting the grid size to the precision required for compound identification, while discarding empty regions and maintaining computational feasibility.

Many peak detection methods rely on a data reduction step (centroiding), in which profile data are collapsed into single representative points (centroids) per peak [6]. While this reduces data volume, it can also distort the signal and discard information that may be valuable for downstream analyses [7]. Choosing centroiding parameters is particularly problematic in untargeted studies where compound classes are not known in advance. Our approach avoids this step by working directly on the raw data, thus preserving full measurement precision.

Prior to decomposition, we apply a simple noise filter by discarding measurements below a fixed intensity threshold (set to the 0.1-quantile of measured intensities in our experiments). This reduces data volume while retaining the informative signal.

### Algorithm

The method consists of two main steps (Figure 2): (i) hierarchical decomposition of the raw data into resolution-dependent regions of interest (ROIs), and (ii) peak detection within each ROI using image-processing techniques.

**Figure 1.**
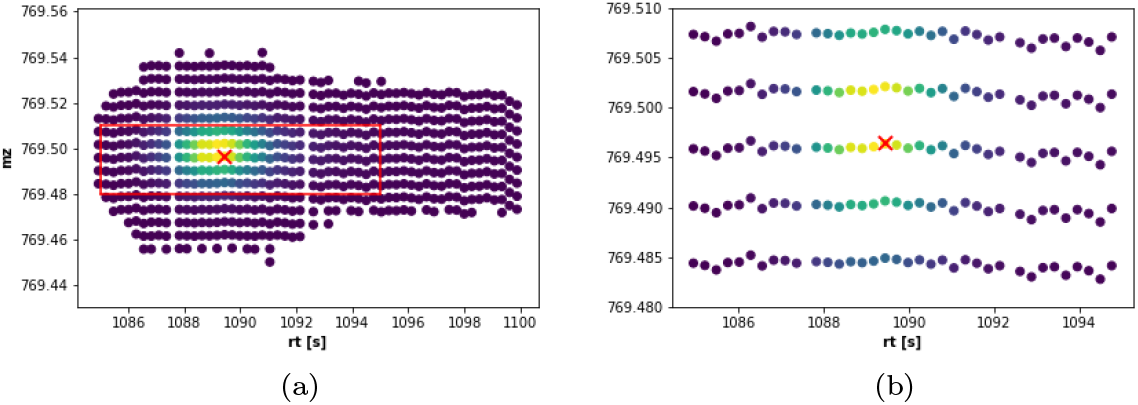
A small region around a peak at low (1a) and high (1b) resolutions, respectively. The dots represent actual measurements and the red cross indicate the local maximum. Intensity (amplitude of the signal) is color coded, with yellow indicating higher values. The red rectangle in (1a) shows the region magnified in (1b)

**Figure 2.**
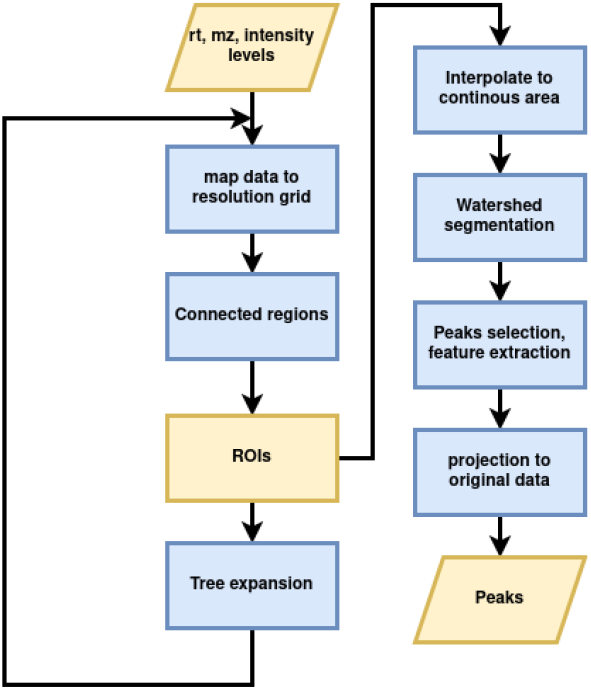
Conceptual workflow of the MRH algorithm. The method consists of two main steps: (i) hierarchical decomposition of the raw data into resolution-dependent regions of interest (ROIs), and (ii) peak detection within each ROI using image-processing techniques. This design preserves measurement precision, enables scalable processing, and allows subsequent evaluation of peak quality using the ambiguity score.

Hierarchical decomposition. The raw data are first projected onto a regular grid defined by a chosen resolution *ρ*. A point *x* is mapped to *x*_*ρ*_ = ⌈*ρx*⌉, and all measurements within a *ρ*× *ρ* bin are aggregated by their maximum intensity:

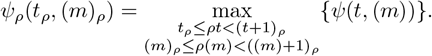

Connected components of non-zero bins are identified using 4-neighbor connectivity. A component is retained as an ROI if it contains at least *n*_min_ = 50 points with non-constant retention time values. Repeating this process across multiple resolutions yields a tree-like decomposition, where coarser ROIs split into finer ones at higher resolutions.

For each ROI, raw measurements within its bounding box and a binary mask are stored, allowing efficient retrieval at any resolution.

Peak detection. Each ROI is treated as a single-channel image. After 2*D* interpolation and smoothing with a 5× 5 rectangular kernel, peaks are identified using a watershed algorithm [10], with markers initialized by the H-maxima transform [11]. Each peak is associated with a support region, from which attributes such as point count, mean, and variance are computed. Peak positions can either be taken from the watershed maxima or back-projected to the closest raw data point for full precision.

Parameterization. The main parameters are the set of explored resolutions *ρ* and the maximum allowed gap when forming ROIs. High resolutions improve precision but increase computational cost, while larger gaps tolerate missing points but risk merging peaks.

### Ambiguity score

As an indirect measure of peak significance, we introduce a score *Q* characterizing the quality of a detected peak. Since its purpose is to flag potentially ambiguous detections, we call it the *ambiguity score*, with values in [0, 1] where higher values indicate more problematic peaks:

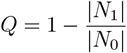

Where

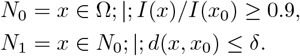

Here, *N*_0_ denotes the set of all data points in the connected area Ω with intensities at least 90% of the peak intensity *I*(*x*_0_). *N*_1_ is the subset of *N*_0_ located within a ball of radius *δ* around the peak apex *x*_0_. Thus, the ambiguity score *Q* reflects the fraction of high-intensity points that lie in the same ROI as *x*_0_ but are farther away than *δ*, potentially indicating overlapping or unresolved peaks. The tolerance *δ* depends on both retention time (RT) and mass-to-charge ratio (*m/z*) and was set equal to the allowed neighborhood gap.

Figure 3 illustrates three representative cases. A well-isolated peak yields *Q* = 0.0 (Figure 3a), while more ambiguous cases include band-like regions (*Q* = 0.758, Figure 3b) and regions containing potentially multiple overlapping peaks (*Q* = 0.636, Figure 3c). A visual explanation of the quantities used in the score definition is provided in Supplemental Figure S1.

**Figure 3.**
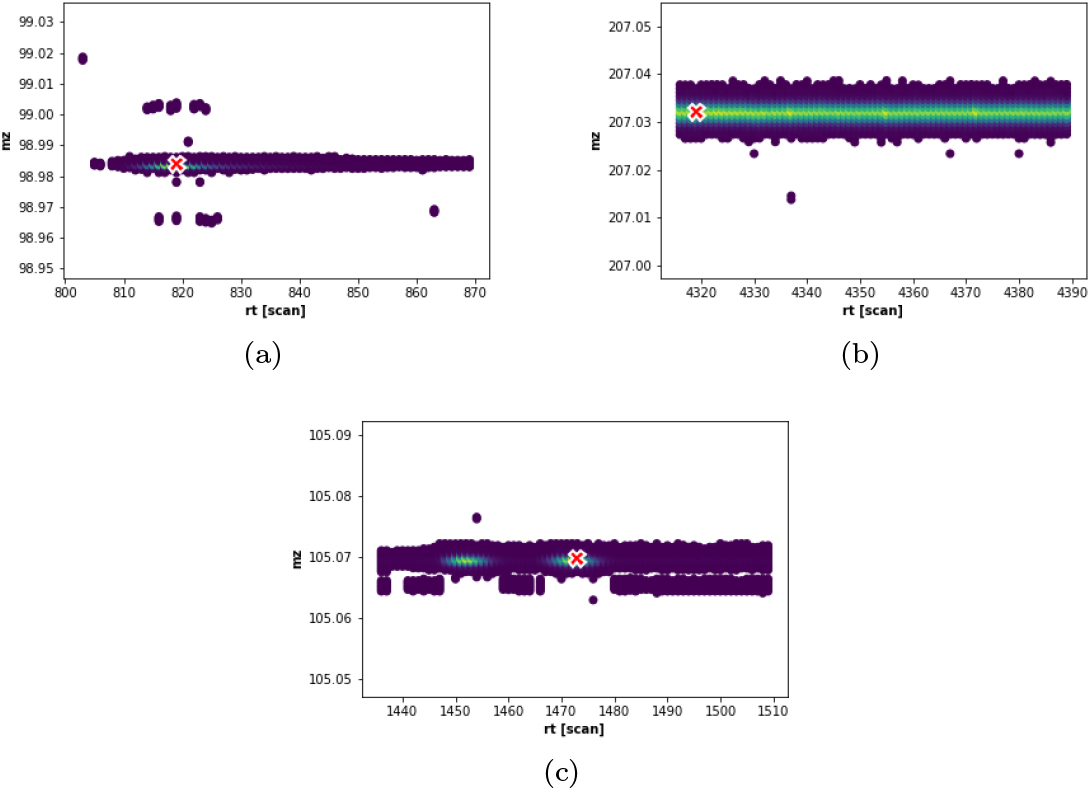
Examples of detected peaks with different ambiguity scores. (a) Well-isolated peak with *Q* = 0.0. Detection in a band-like region (*Q* = 0.758). (c) Region containing a potentially overlapping peak (*Q* = 0.636).

### Test data

To evaluate the proposed method and compare its performance with standard peak detectors, we generated an in-house dataset consisting of mixtures of flame retardants at known proportions and predefined concentrations. The dataset is derived from the INTERFLAB experiment described in [12]. Measurements were performed in profile mode using a Thermo Fisher Scientific Q Exactive GC Orbitrap GC-MS/MS system.

To construct the ground truth list of peaks, the predicted peak positions were matched against the raw data, and the closest local maxima were designated as true peaks. This adjustment ensured alignment of predicted and observed values across samples. An example of such a peak within a small region of the RT/MZ space is shown in Figure 1. In some cases—particularly at lower concentrations—no raw measurements could be associated with certain predicted peaks. These instances were marked as “no data.” Further details of the dataset generation and curation are provided in the Supplemental Materials (S2).

### Performance assessment

Each detector’s output was compared against a manually curated ground truth list. A detected peak was considered a true positive if its retention time was within a tolerance of Δ*RT* = 0.02 s and its mass-to-charge ratio within Δ*MZ* = 0.001 of a ground truth peak. For each ground truth entry, we distinguished between unique detections and multiple detections.

A comprehensive assessment of false positives was not feasible, as the biological samples may contain unannotated or contaminating compounds. From the perspective of a peak detector, however, any peak not present in the ground truth list is treated as unmatched. To account for this, we used precision as a performance metric, reflecting the proportion of matched detections relative to all detections. Combined with recall (the proportion of ground truth peaks successfully detected), this allowed us to calculate the F1 score, which served as our primary measure of overall accuracy. The F1 score provides a balanced evaluation of a tool’s ability to maximize correct detections while minimizing spurious ones.

We compared MRH against several widely used peak detectors: apLCMS, centWave, and gridmass (with both high- and low-tolerance settings). In the gridmass_H setting, the default *m/z* tolerance of 0.1 was applied, whereas in gridmass_L we adjusted the tolerance to a stricter 0.001, the highest feasible resolution. Detailed parameter settings for all methods are provided in the Supplemental Materials (S3).

## Results

We evaluated the performance of MRH against four widely used peak detection tools—apLCMS, centWave, gridmass_H, and gridmass_L — using the benchmark dataset described in Section.

### Overall detection performance

The primary metric for comparison was the F1 score, which balances recall and precision. As shown in Figure 4, MRH consistently outperformed all other methods across the full range of concentrations. The advantage was most pronounced at low concentrations, where competing methods exhibited sharp declines in sensitivity. Even at high concentrations, MRH remained among the best-performing methods, achieving stable and superior F1 scores. Notably, centWave occasionally reached high F1 values at specific concentrations, but its performance was highly variable and inconsistent across replicates.

**Figure 4.**
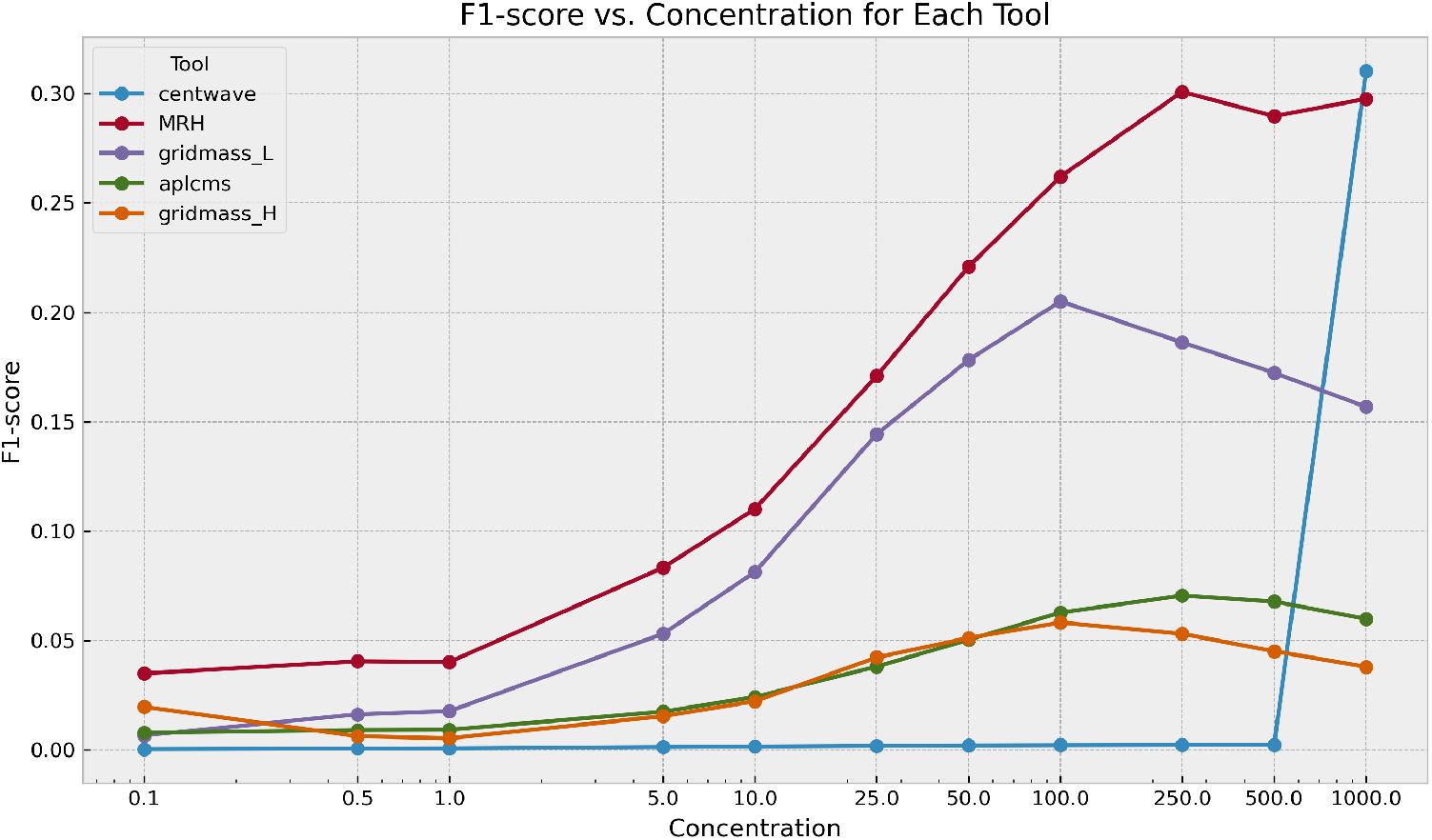
F1 score as a function of concentration for MRH and four competing peak detection tools. MRH achieves consistently higher scores across a wide range of concentrations, with a clear advantage at lower levels.

### Ambiguity analysis with MRH scores

To further characterize detection quality, we analyzed the distribution of MRH ambiguity scores across all detected peaks (Figure 5) and restricted to matched peaks only (Figure 6). Across all concentrations, the majority of peaks exhibited low ambiguity, with mean scores between 0.83 and 0.86. For matched peaks, scores were slightly lower, reflecting the reduced uncertainty in confidently detected peaks. Interestingly, ambiguity tended to increase at the highest concentrations, likely due to denser spectral regions with partially overlapping features.

**Figure 5.**
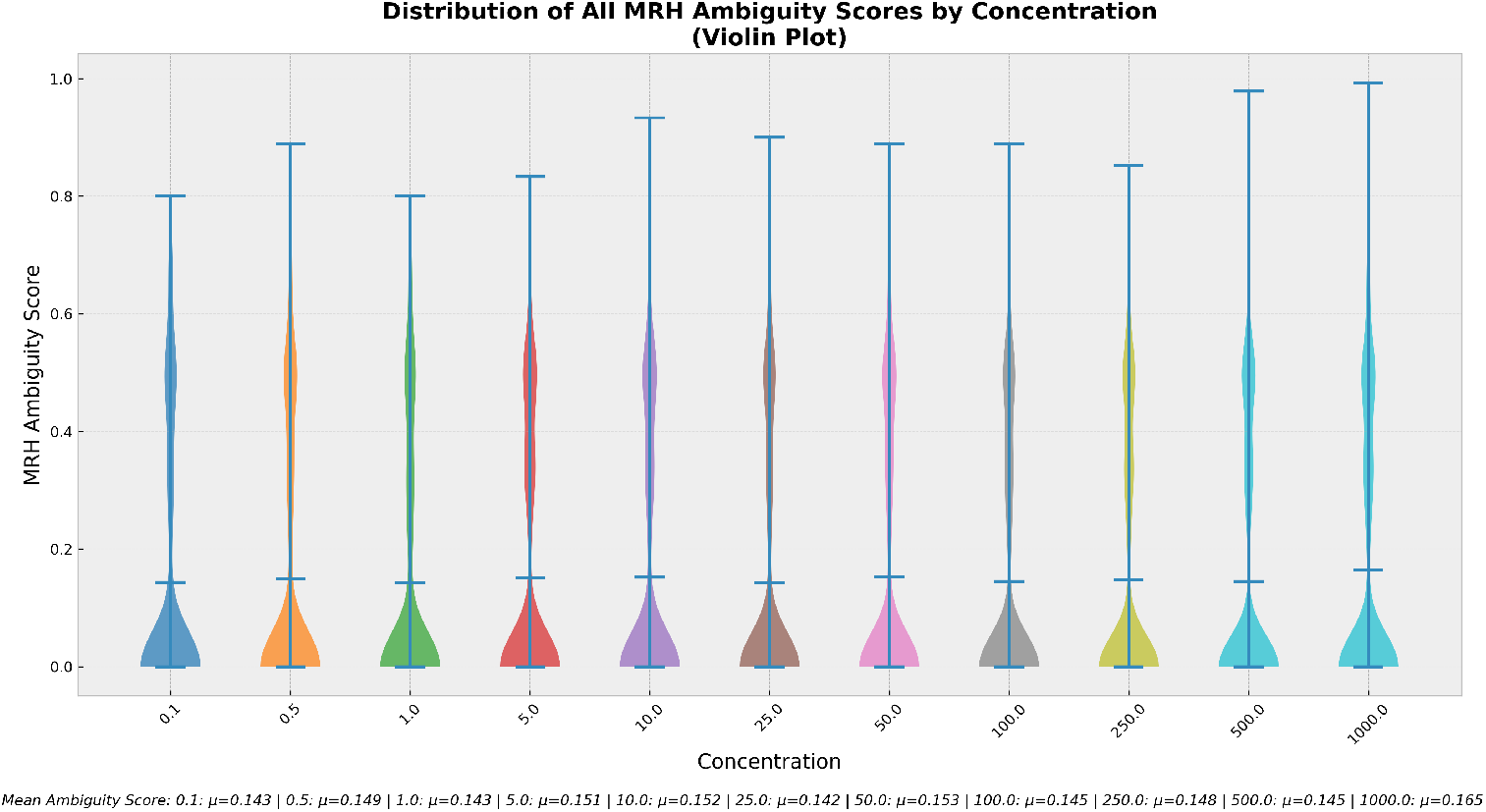
Distribution of MRH ambiguity scores across all detected peaks as a function of concentration. Mean values are indicated for each condition.

**Figure 6.**
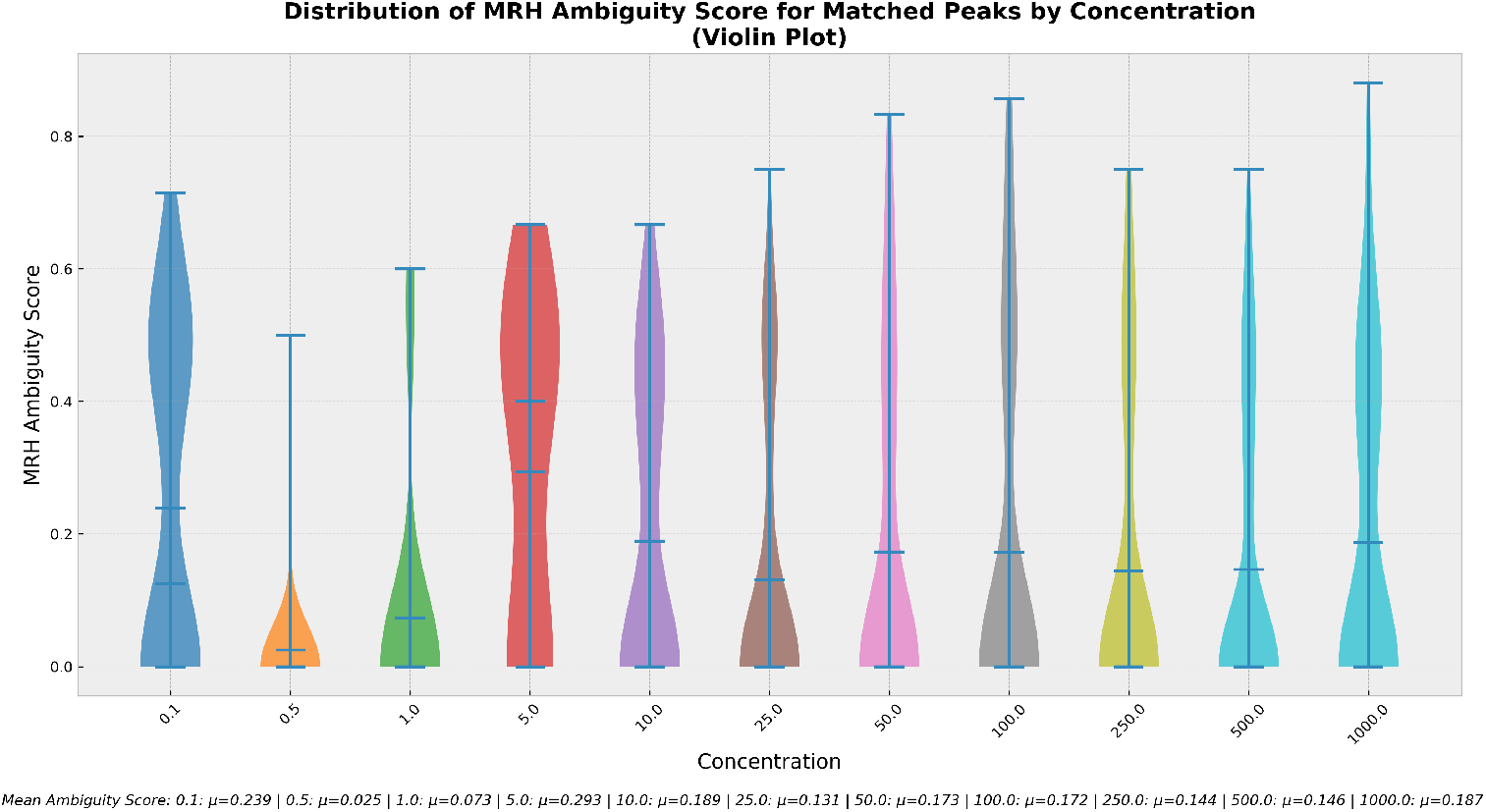
Distribution of MRH ambiguity scores restricted to matched peaks only. Compared to all detections, matched peaks exhibit lower ambiguity, indicating higher confidence in peak assignment.

### Localization accuracy at high concentration

To assess localization accuracy, we compared the differences in retention time and *m/z* between detected and ground-truth peaks at 1000 ng/ml. Contour density plots (Figure 7) show that MRH provided highly accurate localization, with minimal deviations in both dimensions. In contrast, apLCMS and centWave displayed broader error distributions, particularly in the retention time domain, while gridmass variants showed intermediate performance.

**Figure 7.**
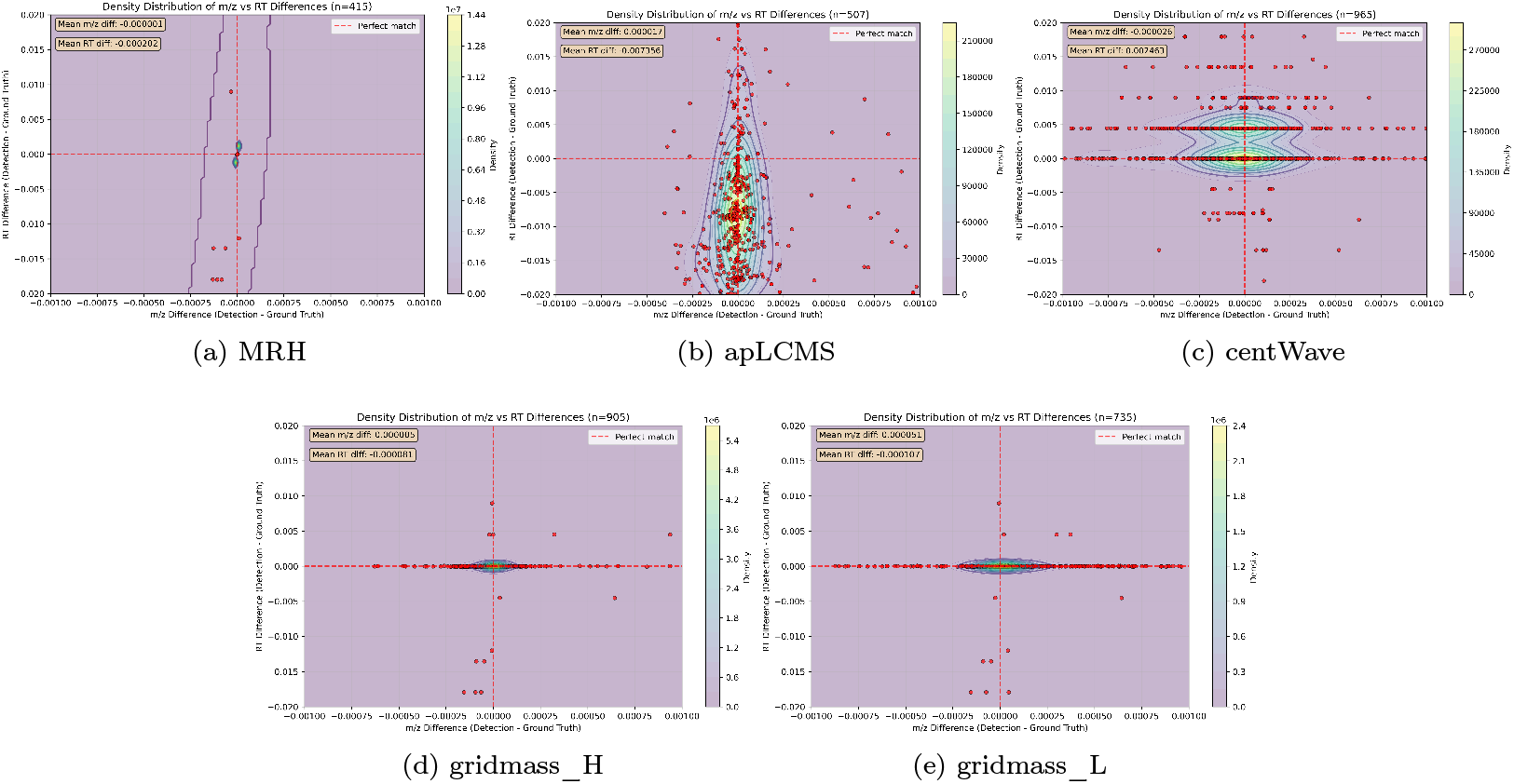
Density contour plots of *m/z* and retention time differences between detected and ground-truth peaks at 1000 ng/ml for MRH and four competing tools. MRH achieves the most accurate localization, with tight clustering around zero error.

### Summary

Taken together, these results demonstrate that MRH delivers superior peak detection performance compared to established methods. It achieves the highest F1 scores across concentrations, produces low-ambiguity detections, and provides accurate peak localization. These findings highlight MRH as a robust and reliable tool for high-resolution mass spectrometry data analysis.

## Discussion and Conclusions

We have introduced MRH, a new approach for peak detection in high-resolution mass spectrometry data, based on hierarchical decomposition of the raw signal into regions of interest followed by image-inspired feature detection. This design addresses several limitations of existing methods. Unlike centroid-based approaches, which irreversibly reduce the data and often introduce distortions, MRH preserves the original raw measurements throughout the analysis. By structuring the data hierarchically, MRH also enables resolution-aware partitioning, efficient traversal of the (RT, *m/z*) space, and scalable parallelization.

Our results demonstrate that MRH delivers superior detection accuracy compared to widely used alternatives such as apLCMS, centWave, and gridmass. Across concentrations, MRH consistently achieved the highest F1 scores, particularly excelling at lower concentrations where other methods struggled. This highlights the robustness of MRH in scenarios where sensitivity is critical. Localization accuracy was also improved, with MRH yielding tight error distributions in both retention time and *m/z* domains.

An additional contribution of MRH is the introduction of the ambiguity score, a simple yet informative measure that quantifies the confidence of each detection. Peaks with low scores represent unambiguous features, whereas higher scores indicate overlapping or less distinct signals. This provides users with a practical means of assessing result quality and adapting analysis strategies, for instance by adjusting resolution or filtering ambiguous detections.

While promising, our study has some limitations. The evaluation was performed on a controlled benchmark dataset with defined ground truth peaks; performance on more complex real-world samples remains to be assessed. Moreover, parameterization of competing methods was kept consistent with standard recommendations, but alternative tuning might affect relative outcomes. Finally, our proof-of-concept implementation does not yet exploit the inherent parallelism of the hierarchical decomposition, leaving room for significant computational optimization.

In summary, MRH combines methodological rigor with practical performance gains. It achieves higher F1 scores, better localization accuracy, and provides a novel measure of detection confidence, all while preserving the richness of raw mass spectrometry data. Together, these properties establish MRH as a robust and reliable peak detection framework with clear potential for integration into large-scale metabolomics and other high-throughput mass spectrometry workflows.

We make available both a proof-of-concept implementation of the described method and the data used for testing it to support further comparison and developments.

## Acknowledgements

This work has been funded by grant SALVAGE (CZ.02.01.01/00/22_008/0004644) from the Programme Johannes Amos Comenius under the Ministry of Education, Youth and Sports of the Czech Republic – Cofunded by the European Union. Authors thanks to Research Infrastructure RECETOX RI (No LM2023069) financed by the Ministry of Education, Youth and Sports, and Operational Programme Research, Development and Innovation - project CETOCOEN EXCELLENCE (No CZ.02.1.01/0.0/0.0/17_043/0009632) for supportive background. Computational resources were provided by the e-INFRA CZ project (ID:90254), supported by the Ministry of Education, Youth and Sports of the Czech Republic. This work was also supported from the European Union’s Horizon 2020 research and innovation program under grant agreement No 857560 (CETOCOEN Excellence). This publication reflects only the author’s view, and the European Commission is not responsible for any use that may be made of the information it contains.

## Supporting information

- S1: Peak ambiguity measure: defining parameters
- S2: Annotation tables
- S3: Detectors settings

