## Supplement for "A Multiresolution Hierarchical Approach to Peak Picking for High Resolution Mass Spectrometry Data Analysis"

### S1: Peak ambiguity measure: defining parameters

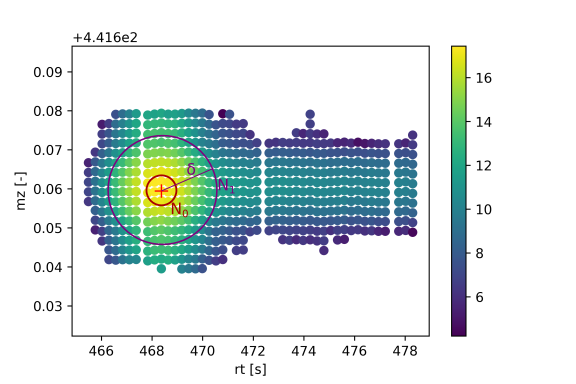

### S2: Annotation tables

Provided annotation table was subsequently edited to be fitted for each concentration. We provide, for each identifiable ion in the raw data, the corrected mz and rt. The predicted positions of the peaks were matched against the raw data and the closest matching local maxima were deemed the true positions of the peaks. This step was done to fit the predicted positions to match actual values in each sample. All corrected values are available at zenodo.

### S3: Detectors settings

For method gridmass_v2 the settings were adjusted to 0.001 for the maximum M/Z tollerance. This settings was choosen as the maximum feasible for the testing machine to process.

| **Detector** | **Parameters** |
| --- | --- |
| Centwave  version xcms 4.8.0 | ppm = 20,  peakwidth = c(5, 60),  snthresh = 10,  prefilter = c(3, 100),  mzdiff = -0.001  integrate = 1L,  mzdiff = -0.001,  fitgauss = FALSE, |
| apLCMS  version 6.6.6 | bandwidth=0.5,  min.bw=NA,  max.bw=NA,  sd.cut=c(1,60),  sigma.ratio.lim=c(0.2, 5),  shape.model="bi-Gaussian",  estim.method="moment",  do.plot=TRUE,  power=2,  component.eliminate=0.01,  BIC.factor=2 |
| GridMass_L  version mzmine 4.7.8 | minimum_height = 20,  mz_tolerance = 0.1,  min_max_width_time = (0.1, 3.0),  smoothing_time = 0.05,  smoothing_mz = 0.05,  false_intensity_ratio = 0.5 |
| GridMass_H  version mzmine 4.7.8 | minimum_height = 20,  mz_tolerance = 0.001,  min_max_width_time = (0.1, 3.0),  smoothing_time = 0.05,  smoothing_mz = 0.05,  false_intensity_ratio = 0.5 |
